# E-InfertilityTest: An Explainable AI Framework for Male Infertility Assessment

**DOI:** 10.64898/2026.05.21.726746

**Authors:** Gourab Das, Byapti Ghosh, Zhumur Ghosh

## Abstract

Male infertility has emerged as a significant concern in modern society, with genetic defects as one of the major underlying cause behind it. This impairment negatively impacts sperm motility and morphology, leading to conditions such as Asthenozoospermia (reduced sperm motility), Teratozoospermia (abnormal sperm morphology) and sometimes Asthenoteratozoospermia (both motility and morphology defects). Assisted reproductive technologies (ART), such as in-vitro fertilization (IVF), offer a potential solution for such cases but with a low success rate. Classical semen analysis provides only a phenotypic snapshot without revealing the fertilizing potential of the sperms. Hence, in order to screen the functional sperm population as well as to get a deeper insight into the reasons underlying the aberrant sperm population, it is important to study their genetic profile. In this work, we have performed a meta analysis of the transcriptomic data of infertile sperms from Asthenozoospermia and Teratozoospermia patients with that from fertile sperms of normal individuals. Thereafter we have screened a signature gene set which has been used to develop a prediction model named **Explainable Infertility Test (E-InfertilityTest)** to classify between fertile versus infertile sperm at the preliminary level. For each prediction, it will also provide the set of genes which are playing a dominant role towards such prediction. Thus, it will provide patient specific dominant gene expression profile responsible for the aberration. This work warrants validation experiments in future to substantiate the model’s performance in a clinical setting. User can access the tool named E-InfertilityTest as a standalone version on GitHub.

**Github Link:** https://github.com/zglabDIB/einfertility.git

## INTRODUCTION

Infertility has emerged as a significant global health concern, affecting approximately 15% couples worldwide, with male factors contributing to 20-30% of these cases [1]. Among the abnormalities assessed in standard semen analysis, defects in sperm motility (asthenozoospermia) and morphology (teratozoospermia) constitute important, often under-recognized causes of male subfertility. Assisted reproductive technologies (ART), such as in-vitro fertilization (IVF), offer a potential solution for such cases, but the overall success rate remains low [2]. Efforts to enhance the success of IVF have focused on pre-screening of sperms based on various molecular, morphological and physiological parameters [3, 4]. Various clinically used sperm preparation techniques to separate functional sperms from semen plasma also play a significant role in the process [5]. However, classical semen analysis provides only a phenotypic snapshot without revealing the fertilizing potential of the sperms in terms of their underlying genetic profile which is not only important in detecting the functional sperm population but also reveal the genetic reasons underlying the aberrant sperm population. Thus, in such cases of fertility defects, there is a growing need to get a deeper insight into the genetic profile underlying sperm defects, which will further help to obtain an enriched fertile sperm population, thereby increasing the success rate of IVF. Male infertility is commonly assessed through clinical parameters such as sperm count, motility and morphology [6]. Motility is necessary for sperm to navigate the female reproductive tract and reach the oocyte, while morphology reflects the structural integrity required for normal sperm-egg interaction, acrosome reaction and chromatin integrity [7, 8]. Abnormalities in either attribute have been associated with reduced fertility but the molecular etiology often remains elusive, especially in idiopathic male infertility [9]. By focusing on asthenozoospermia and teratozoospermia, we target a subset of male infertility cases where functional impairment (rather than just sperm count) may be derived from underlying genetic profile.

The initial step in diagnosing male infertility involves conventional semen analysis, which assesses spermatogenesis and glandular secretory activity [10]; while serum hormonal levels, including LH, FSH, total testosterone, E2, PRL and T/E2, are also evaluated in clinical testing [11]. Given the correlation between semen analysis results and serum hormone levels, Kobayashi et al. [12] investigated the possibility of using machine learning (ML) to predict male infertility based solely on serum hormone levels. Arroyo et al. [13] has published very informative review regarding the implication of ML based artificial neural network in order to predict male infertility. Based on their findings, majority of prediction models are focused on Azoospermia (absence of sperm in the ejaculate). Several studies have been done on sperm retrieval, IVF outcomes and the influence of environmental and medical factors [13]. Additionally, many prediction models have explored sperm quality and morphology, although their primary conclusions are derived from their videos and images [13]. On the other hand, high-throughput transcriptomic technologies offer a powerful route to uncover gene expression changes associated with sperm defects. Understanding the genetic basis of male infertility is crucial for providing effective solutions to couples seeking assistance. Notably, Bansal et al. [14] identified coding RNAs in sperm, proposing their potential as biomarkers for fertility assessment. Further, Shamsi et al. [15] emphasized the importance of genomic integrity in germ cells as a major factor in fertility. Jenkins et al. [16] performed an experiment using semen samples from 94 men to identify the perturbed cellular pathways corresponding to teratozoospermia and asthenozoospermia. In a more recent study performed by Li et al. [17], metabolomics profiling have been studied to identify distinct metabolites related to similar diseases using ML approaches.

Although there exist some studies involving gene expression profiles to differentiate between normal and asthenozoospermic [18] or teratozoospermic [19] samples, but there lies significant gap regarding development of ML based model to predict the status of a sperm sample by considering gene expression as feature sets. In this work, we have performed a rigorous transcriptomic analysis of the gene expression profile of infertile (asthenozoospermic and teratozoospermic patient sperm sample) versus fertile sperm sample to screen a signature gene set. This set have been further filtered and used to develop a prediction model named Explainable Infertility Test abbreviated as **E-InfertilityTest** to classify between fertile and infertile sperm. In order to mitigate the issue of data dimension, we have deliberately adopted the simplest linear classifier in forms of Logistic Regression as it constructs a linear decision boundary with lower representational complexity, which is generally less prone to overfitting in small-sample settings. Moreover, we have built a novel feature selection algorithm which recursively includes the gene expression features in an iterative manner to improve the efficacy of our model. Finally, we have deployed an Explainable AI framework [20], to further identify the set of genes playing a dominant role towards such classification. Thus, it will provide patient specific dominant set of genes responsible for the aberration.

User can access standalone version of E-InfertilityTest from ***https://github.com/zglabDIB/einfertility.git***

## MATERIAL AND METHODS

The detailed workflow associated with **E-InfertilityTest**, depicted in ***Figure 1***, can be segregated into two phases. The initial phase deals with a rigorous bioinformatics analysis to elucidate the significantly dysregulated genes and their associated pathways corresponding to infertile sperm samples with respect to the normal counterparts. We have further used this refined gene set to develop an ML based fertility prediction model. Moreover, an Explainable AI framework has been developed to identify sample specific dominant set of genes for each prediction.

**Figure 1:**
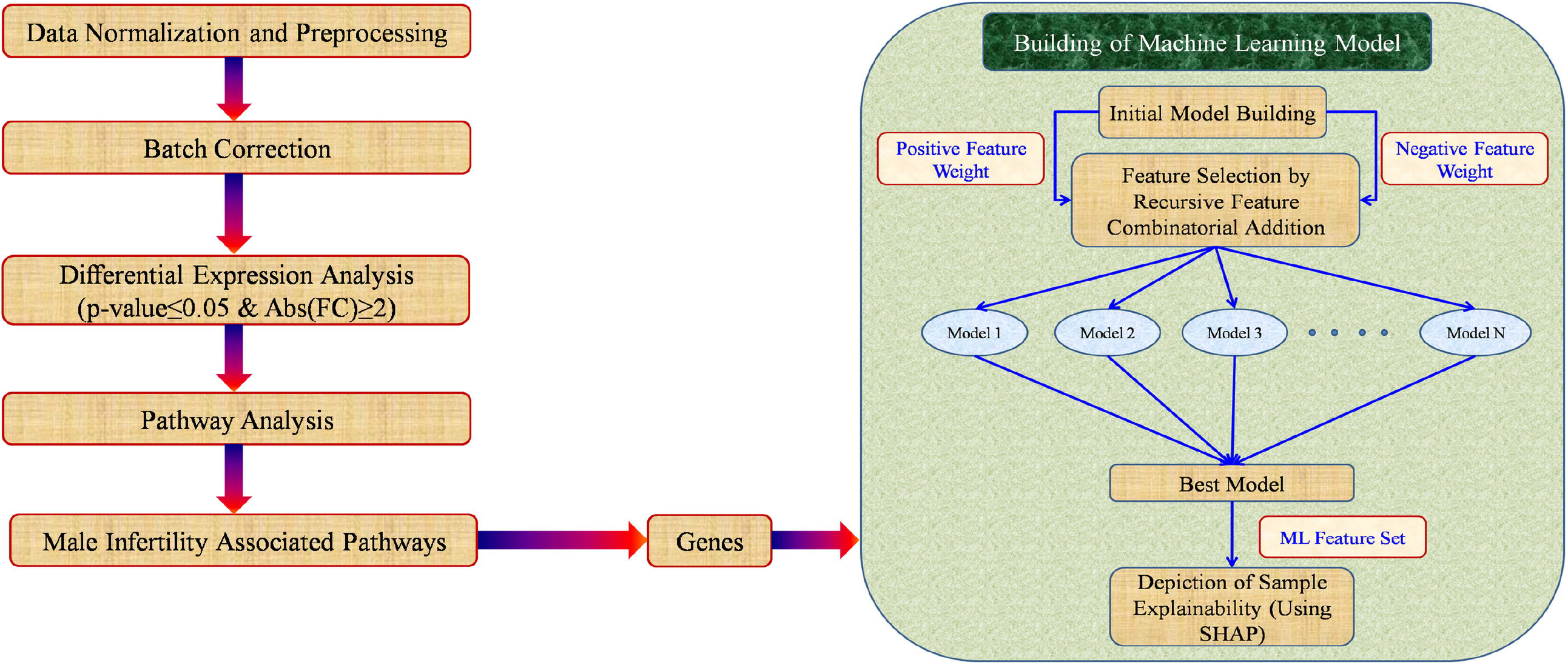
Workflow of the entire work. Flow diagram depicts the various steps involved towards the development of E-InfertilityTest which includes Bioinformatics analysis and the construction of Machine Learning model

### Input Dataset

We have downloaded the microarray gene expression dataset from NCBI GEO databases (https://www.ncbi.nlm.nih.gov/gds) corresponding to human sperm samples from patients having fertility defects viz. asthenozoospermia and teratozoospermia as well as that from normal counterparts (depicted in ***Table 1 and Table 2***).

**Table 1:** Training dataset corresponding to fertile and infertile sperm samples.

| Infertility Type | Experiment<br>Accession Number | Sample Accession Number | Platform information | # of<br>infertile<br>samples | # of fertile<br>samples |
| --- | --- | --- | --- | --- | --- |
| Asthenozoospermia | GSE34514 | GSM850437 | Microarray (Affymetrix Human<br>Genome U133 Plus 2.0) | 14 | 19 |
|  |  | GSM850432 |  |  |  |
|  | GSE160749 | GSM4878471 | Microarray(Affymetrix Human<br>Gene 2.1 ST Array) |  |  |
|  |  | GSM4878481 |  |  |  |
|  |  | GSM4878482 |  |  |  |
|  |  | GSM4878483 |  |  |  |
|  |  | GSM4878484 |  |  |  |
|  |  | GSM4878485 |  |  |  |
| Teratozoospermia | GSE6872 | GSM158465 | Microarray (Affymetrix Human<br>Genome U133 Plus 2.0) |  |  |
|  |  | GSM158466 |  |  |  |
|  |  | GSM158467 |  |  |  |
|  |  | GSM158468 |  |  |  |
|  |  | GSM158469 |  |  |  |
|  |  | GSM158470 |  |  |  |
|  |  | GSM158471 |  |  |  |
|  |  | GSM158472 |  |  |  |
|  |  | GSM158473 |  |  |  |
|  |  | GSM158476 |  |  |  |
|  |  | GSM158477 |  |  |  |
|  |  | GSM158478 |  |  |  |
|  |  | GSM158479 |  |  |  |
|  |  | GSM158480 |  |  |  |
|  |  | GSM158481 |  |  |  |
|  | GSE6967 | GSM160620 | Microarray (Sentrix Human-6<br>Expression BeadChip) |  |  |
|  |  | GSM160621 |  |  |  |
|  |  | GSM160622 |  |  |  |
|  |  | GSM160623 |  |  |  |
|  |  | GSM160624 |  |  |  |
|  |  | GSM160626 |  |  |  |
|  |  | GSM160629 |  |  |  |
|  |  | GSM160630 |  |  |  |
|  |  | GSM160631 |  |  |  |
|  |  | GSM160632 |  |  |  |

**Table 2:**
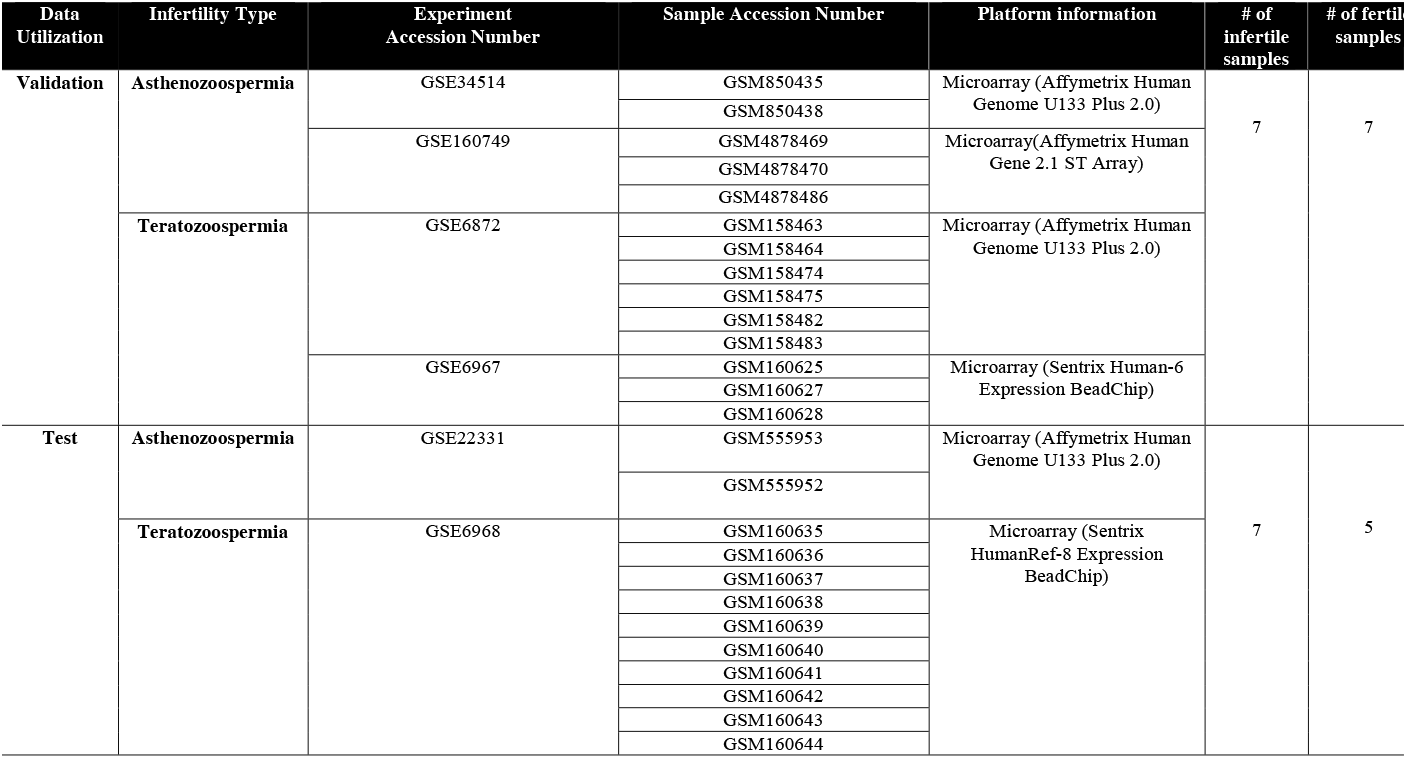
Validation and Test dataset corresponding to fertile and infertile sperm samples.

### Raw data normalization and analysis

The Robust Multi-array Average (RMA) [21] algorithm has been employed for the Affymetrix data and the NormExp Background Correction and Normalization (neqc) [22] have been used to normalize Illumina dataset. Both methods involve background correction followed by quantile normalization, with RMA additionally performing summarization using robust averaging techniques. Batch correction has been done using the Combat [23] tool. Linear Models for Microarray Data (LIMMA) [24], an Empirical Bayes method is utilized to perform differential expression analysis corresponding to the microarray dataset. DEGs were further screened using a p-value cut off of ≤0.05 and Fold Change (FC) cut off of ≥ 2.0.

### Pathway Analysis

The DEGs have been subjected to Ingenuity Pathway Analysis (IPA) (www.qiagenbioinformatics.com/products/ingenuity-pathway-analysis/) [25] for their functional annotation. Subsequently, a rigorous literature survey has been done to screen the pathways involved in male infertility. The genes involved in the screened pathways further serve as the signature gene set corresponding to male fertility defect.

### Machine Learning Model creation and validation

In the next phase of this work, we have developed a prediction model named **Explainable Infertility Test** abbreviated as **E-InfertilityTest** using Logistic Regression [26] based classification and dimensionality reduction based supervised Uniform Manifold and Approximation (UMAP) [27] visualization to classify unknown sperm samples. Moreover, we have used model agnostic SHAP (SHapley Additive exPlanations) [20] tool to incorporate explainability by computing the gene signature specific feature contribution responsible for the fertility status of an unknown sample.

### Initial Model Building

Using the signature gene set identified from male infertility-associated pathways in the previous step, a Logistic Regression model has been constructed. This approach yielded feature importance scores, represented by both positive and negative coefficients or feature weights, which have been used in the feature selection process in subsequent stages.

### Feature Selection by Recursive Feature Combinatorial Addition Method

Corresponding to each of the features having positive coefficient, negative features are combined and added recursively to perform feature selection in cumulative manner, which are subsequently utilized for training procedure using the logistic regression model to extract the prediction metrics upon validation dataset. The main objective is to identify a set of genes from the signature gene set which will serve as an optimized feature set and can classify between fertile and infertile samples. Based on these optimized feature set, the final logistic regression-based classification model and dimensionality reduction-based supervised UMAP visualization model have been developed. The supervised UMAP model provides a visual representation of the status of validation and unknown test dataset(s) with respect to the training dataset.

Deploying Explainable AI framework: In order to compute the contribution of each feature to predict patient specific sample status, we have deployed an Explainable AI framework through SHAP (SHapley Additive exPlanations) package. It yields **Shapley values** generated from cooperative game theory to fairly distribute the payout among players based on their contribution. Here, it computes the contribution of each feature which is calculated by the marginal coalition based averaging across all features.

Corresponding to M number of features of sample x, the φ_i_(x) denotes the feature specific SHAP values where i=1, 2, 3, ………, M. The model’s prediction for sample x, denoted f(x) can be computed using formula given in Equation 1:

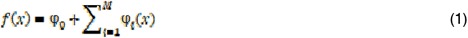

Where 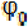 denotes the base or expected value of SHAP explainer calculated over training dataset

For the binary classification problem SHAP extracts log-odds ratio as output. Using the logistic or sigmoid function log-odds ratio can be converted into the prediction probability (p) given as in Equation 2.

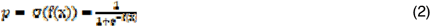

### Software

Both R and Python programming languages have been incorporated in order to create the in-silico analysis based prediction tool. R based libraries are mainly downloaded from Bioconductor (https://www.bioconductor.org/) software packages, whereas Anaconda (https://www.anaconda.com/products/distribution) served for Python based prediction tool. Affymetrix microarray data have been normalized using rma method from affy (https://www.bioconductor.org/packages/release/bioc/html/affy.html) package whereas neqc method from LIMMA (https://bioconductor.org/packages/release/bioc/html/limma.html) package is utilized to perform normalization operation corresponding to illumina microarray data. ComBat method from sva (https://www.bioconductor.org/packages/release/bioc/html/sva.html) package performs the batch correction; the differential expression has been done using the linear modeling from LIMMA package. Fundamental ML model has been created with *Scikit-learn* (https://scikit-learn.org/stable/) [28], whereas UMAP algorithm has been executed using the umap package itself (https://umap-learn.readthedocs.io/en/latest/). The explainable framework has been incorporated using the SHAP (https://shap.readthedocs.io/en/latest/index.html) tool. Finally, we have utilized both ***Matplotlib*** (https://matplotlib.org/) from python and ***ggplot2*** (https://ggplot2.tidyverse.org/) from R to create various plotting.

## RESULTS

### Global gene expression profile of sperm samples with fertility defects

Global gene expression profile have been obtained for infertile (asthenozoospermic and teratozoospermic) and fertile sperm samples. The input dataset serving as the training data and experimental grouping is provided in ***Table 1***. Differential expression analysis has been executed upon 33 training samples (19 fertile and 14 infertile sperm samples). A significant set of 893 DEGs have been obtained with a p-value cut off of ≤ 0.05 and fold change cut off of ≥ 2.0. These DEGs have been subjected to pathway analysis using IPA. Screening of male infertility enriched pathways from these, resulted in 92 pathways involving 297 DEGs. ***Figure 2*** depicts the heatmap which reveals separate clustering of infertile and fertile samples based on the gene expression pattern of these 297 DEGs. The dot plot shown in ***Figure 3*** provides the male infertility related 92 pathways. Hence, the associated 297 genes have been further employed to perform ML based prediction task. The validation and test datasets are provided in ***Table 2***.

**Figure 2:**
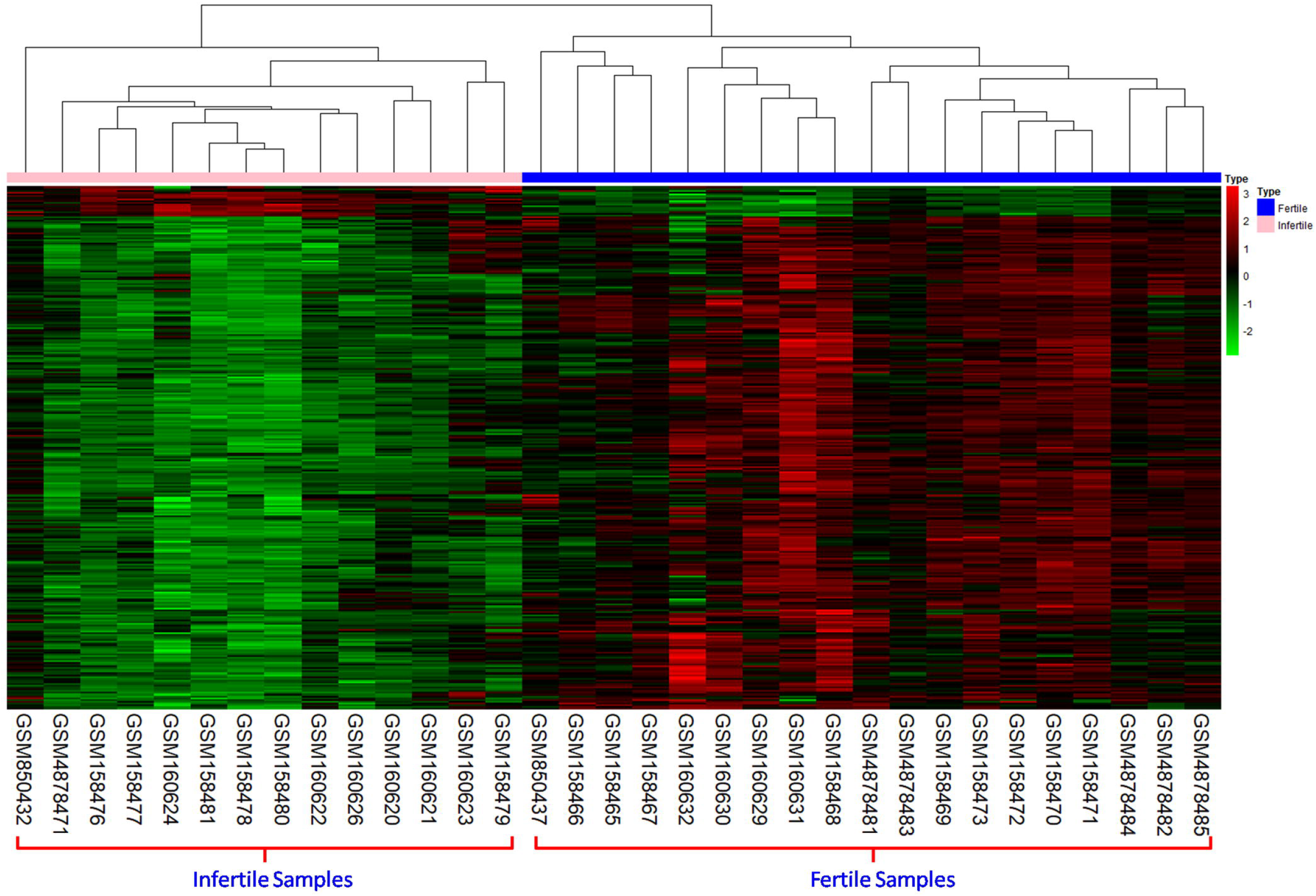
Hierarchical clustering representing global gene expression pattern of the fertile versus infertile sperm samples. Heatmap of 297 DEGs show separate clustering of the fertile and infertile sperm samples

**Figure 3:**
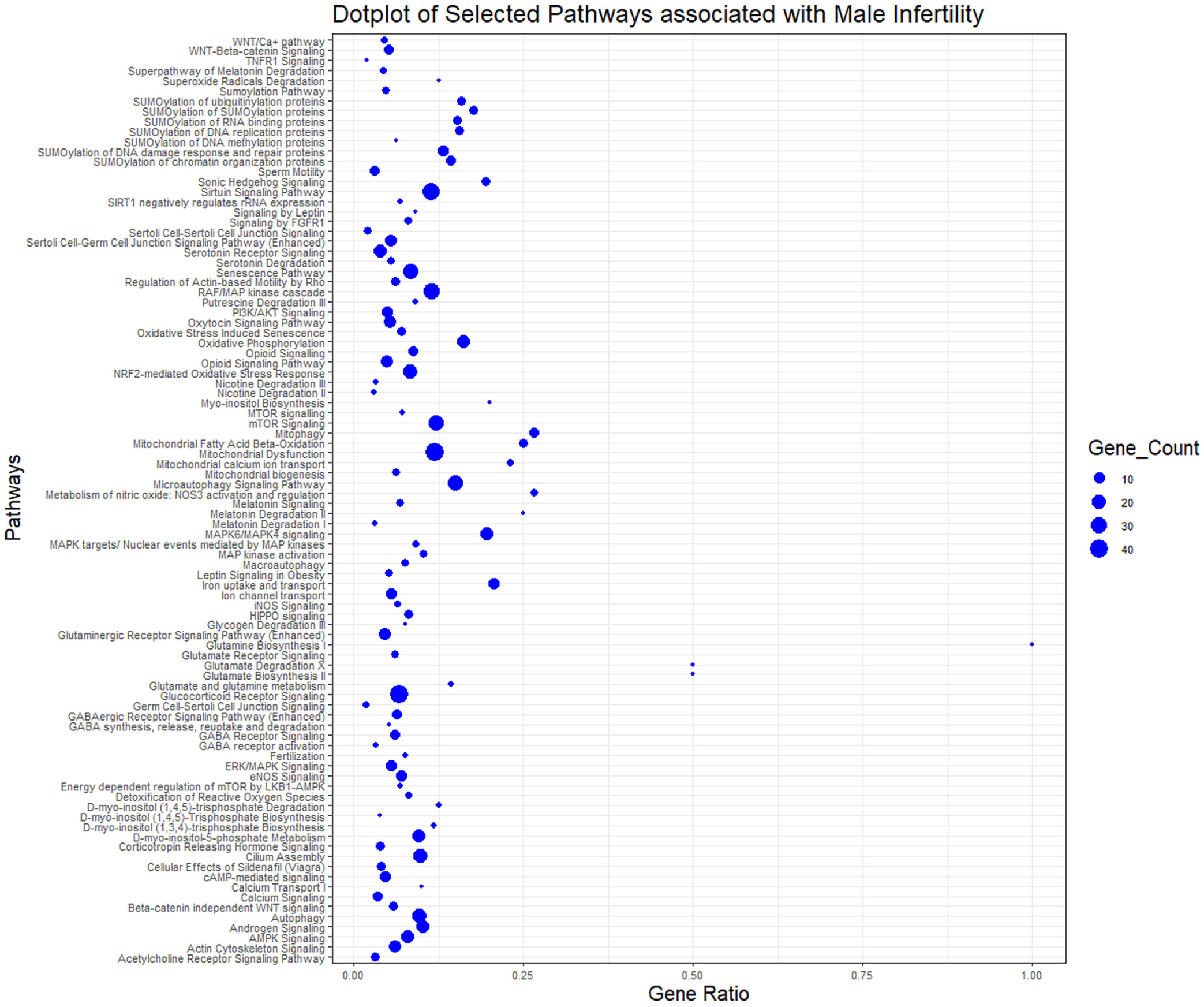
Dotplot showing the pathway specific gene counts associated with male infertility. Dotplot representing 92 pathways involving 297 DEGs elucidated from IPA analysis revealing the male infertility related pathways.

### Implication of ML based Prediction Tools

ML based prediction strategy has been employed to perform the computational correlation operation, where genes associated with the selected pathways have been utilized as feature descriptors during the training process. Two primary operations have been carried out: Logistic Regression-based classification and dimensionality reduction through supervised UMAP visualization. Initially, a preprocessing step in form of normalization has been applied to the training data using StandardScaler, as defined by Scikit-learn. By applying a novel feature selection technique named as the Recursive Feature Combinatorial Addition, the gene set has been narrowed down to 65 genes as the final feature set for training and subsequent prediction tasks. **Supplementary File S1** provides the detailed pathways corresponding to this set of genes. Our logistic regression model has achieved 100% accuracy on both the validation and unknown test data, depicting the significance of the finally selected feature set of 65 genes and their associated pathways, which successfully distinguish fertile samples from infertile ones. ***Table 3*** specifies the prediction probabilities involving both validation and unknown test dataset. ***Figure 4a*** and ***4b*** illustrates the 2D scatter plot generated using the UMAP algorithm, clustering the validation and test samples respectively along with the training samples.

**Table 3:** Prediction Probabilities based on Logistic Regression corresponding to the Validation and Test Data.

| Dataset Type | Sample ID | Prediction Probability |
| --- | --- | --- |
| Validation Dataset | GSM850438_Control | 0.284266 |
|  | GSM850435_Test | 0.750625 |
|  | GSM4878469_Test | 0.999777 |
|  | GSM4878470_Test | 0.999628 |
|  | GSM4878486_Control | 0.002824 |
|  | GSM160625_Control | 0.000191 |
|  | GSM160627_Test | 0.990798 |
|  | GSM160628_Test | 0.999657 |
|  | GSM158463_Control | 0.000351 |
|  | GSM158464_Control | 0.093733 |
|  | GSM158474_Control | 0.002488 |
|  | GSM158475_Control | 0.002419 |
|  | GSM158482_Test | 0.999843 |
|  | GSM158483_Test | 0.999999 |
| Test Dataset | GSM555953_Test | 0.84952 |
|  | GSM160642_Control | 0.075994 |
|  | GSM555952_Control | 0.137013 |
|  | GSM160636_Test | 0.985004 |
|  | GSM160638_Test | 0.534429 |
|  | GSM160640_Test | 0.70353 |
|  | GSM160639_Test | 0.521663 |
|  | GSM160644_Control | 0.120325 |
|  | GSM160641_Control | 0.259073 |
|  | GSM160637_Test | 0.656715 |
|  | GSM160643_Control | 0.047694 |
|  | GSM160635_Test | 0.744664 |

**Figure 4:**
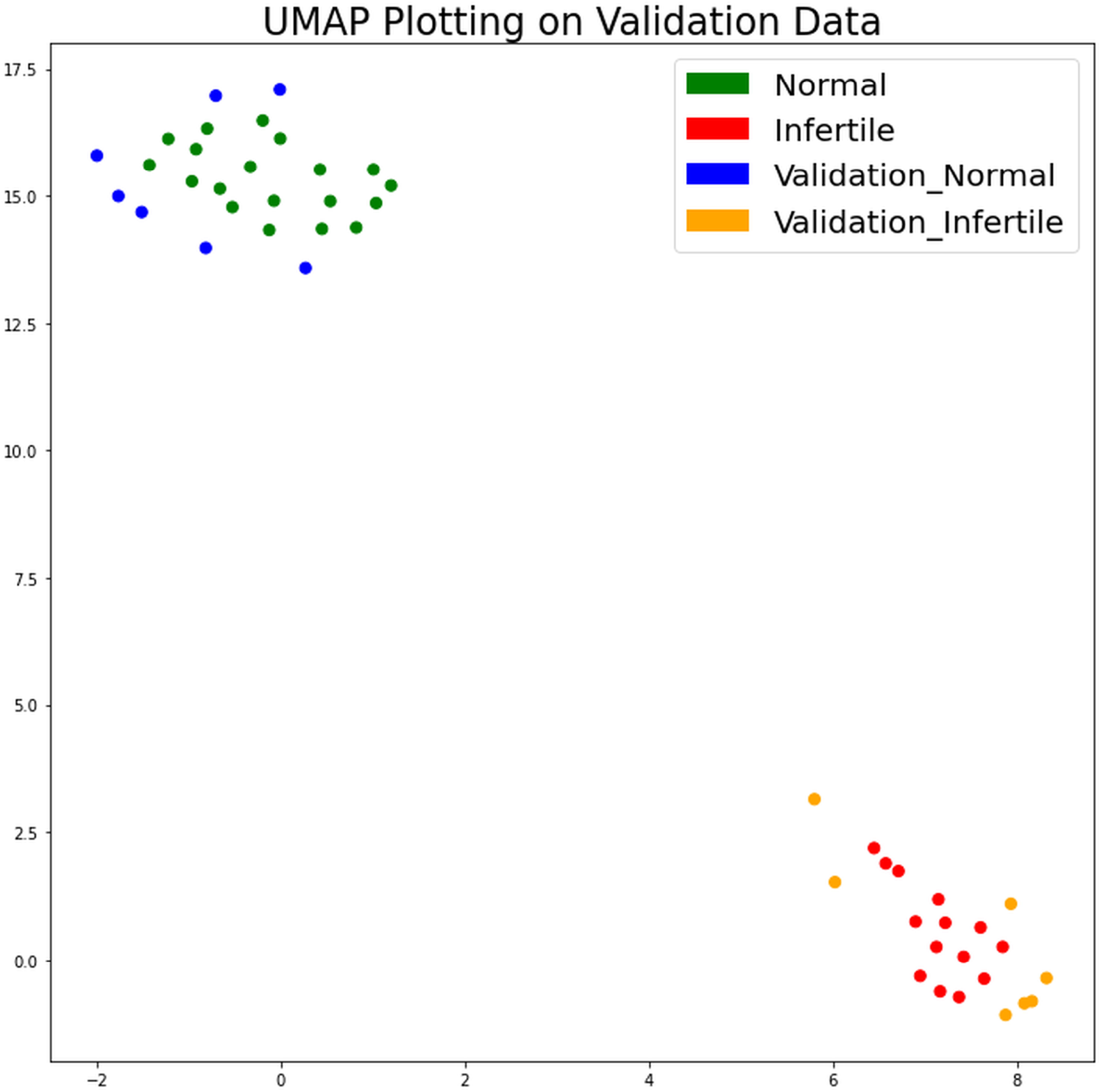
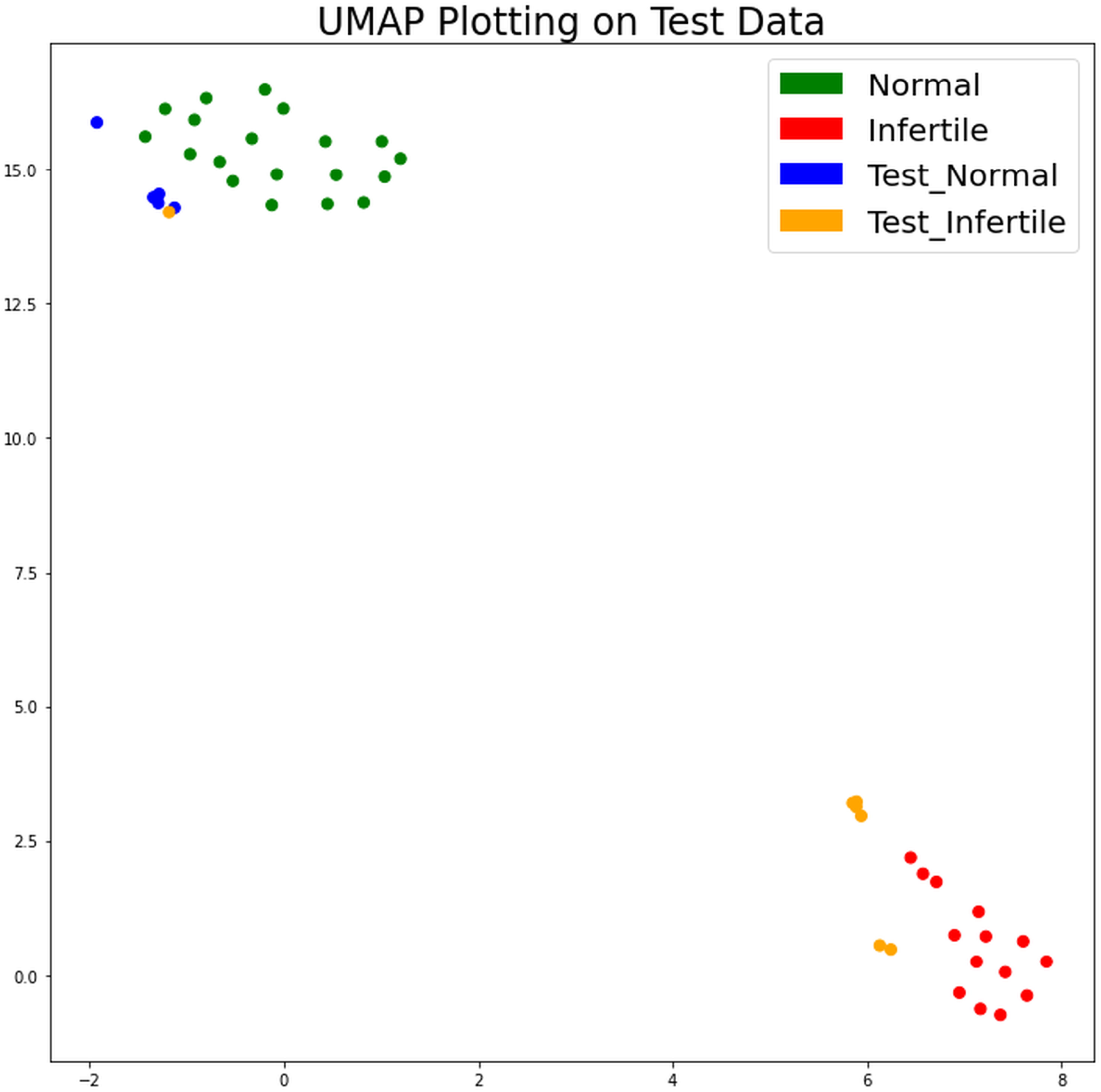
UMAP Visualization of Validation and Test data. 2D Scatter plot representing the cluster of fertile and infertile samples corresponding to (a) Validation and (b) Test samples generated using supervised UMAP algorithm

Despite training on a dataset with 65 features essentially representing 65 dimensions, logistic regression which is a linear classifier flawlessly separates fertile from infertile samples with perfect accuracy. Simultaneously, the UMAP algorithm efficiently projects the entire dataset into a 2-D representation which preserves the underlying structure while simplifying its visualization. Results obtained from logistic regression analysis in ***Table 3*** can be further visualized by UMAP clustering in ***Figure 4***. Among these, the prediction probability for the test data viz. GSM160639_Test, is 0.521 (provided in Table 3) i.e. the sample is almost equally probable to be fertile and infertile. This instigated us to look into the feature specific contribution involved in the prediction outcome for a particular patient specific sample, which will not only provide a deeper insight regarding the reason behind such prediction, but will also assist in personalized therapy.

### Computing feature specific contribution

For an unknown sample, feature specific contribution involved in the prediction is computed using the model agnostic SHAP framework. The force plot generated by SHAP package in ***Figure 5*** depicts the contribution of the genes in decision making. The red and blue arrow indicate the contribution of the genes (from within the feature gene set) which classify the sample to be infertile and fertile respectively. Interestingly, for the sample GSM160639_Test (which is equally probable to be fertile and infertile), the force plot shows almost equal no. of red and blue arrows which means an equal number of genes are playing a dominant role to classify the sample to be fertile and infertile. Hence the probability is around 0.5. On the other hand, force plot for GSM160643_Control sample shows higher number of blue arrows indicating more number of genes which classify the sample to be fertile. For GSM555953_Test sample, higher number of red arrows indicate higher number of those genes which classify the sample to be infertile. ***Figure 6a-6c*** represents the waterfall plot which shows specific set of top 20 most important genes responsible for the prediction outcome in case of GSM160639_Test, GSM160643_Control and GSM555953_Test sample respectively, sorting the set of genes based on their significance.

**Figure 5:**
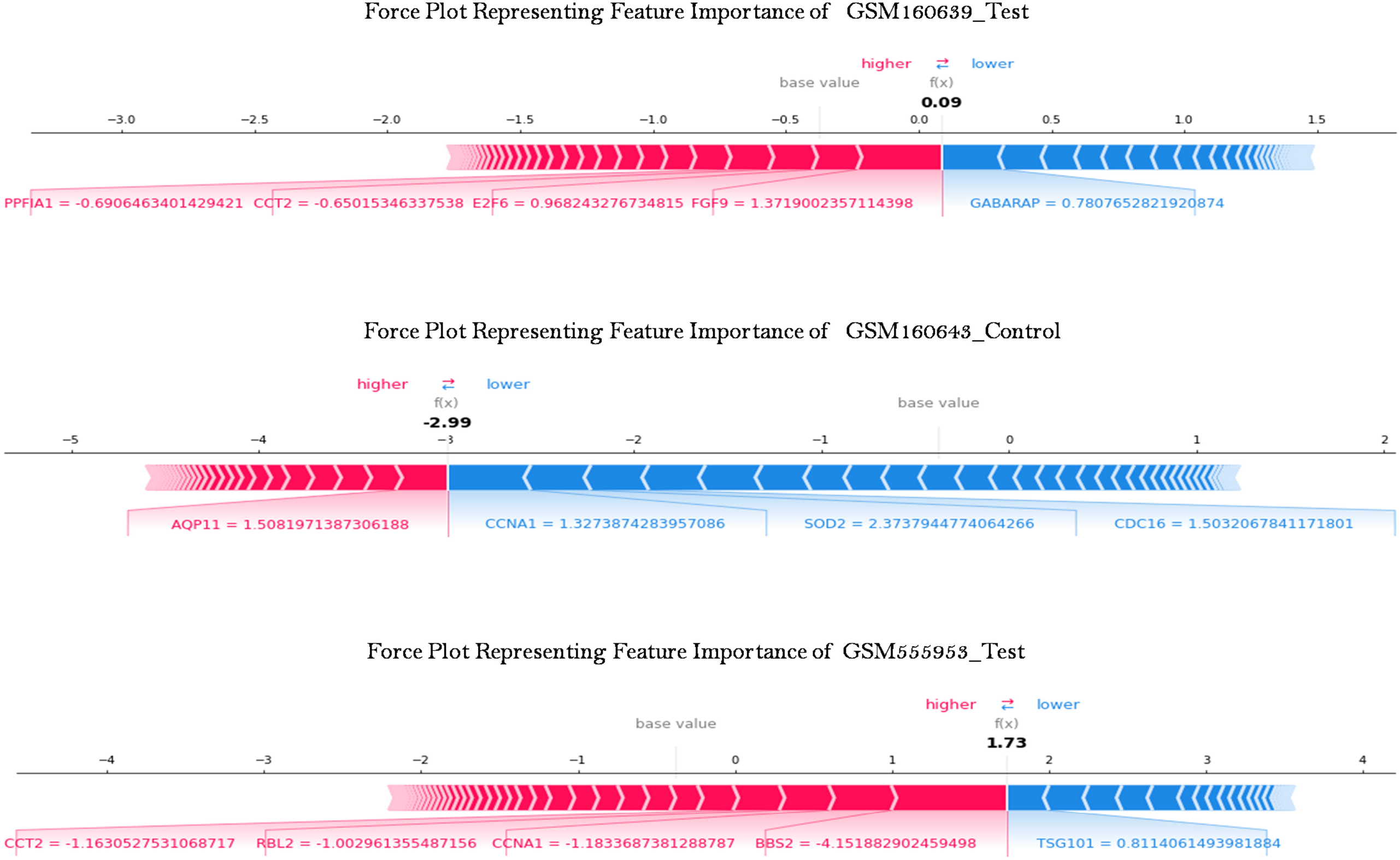
Force Plot summarizing the feature contribution. SHAP generated force plot corresponding to GSM160639_Test, GSM160643_Control and GSM555953_Test samples where red and blue arrow indicate the contribution of the genes (from within the feature gene set) which classify the sample to be infertile and fertile respectively

**Figure 6:**
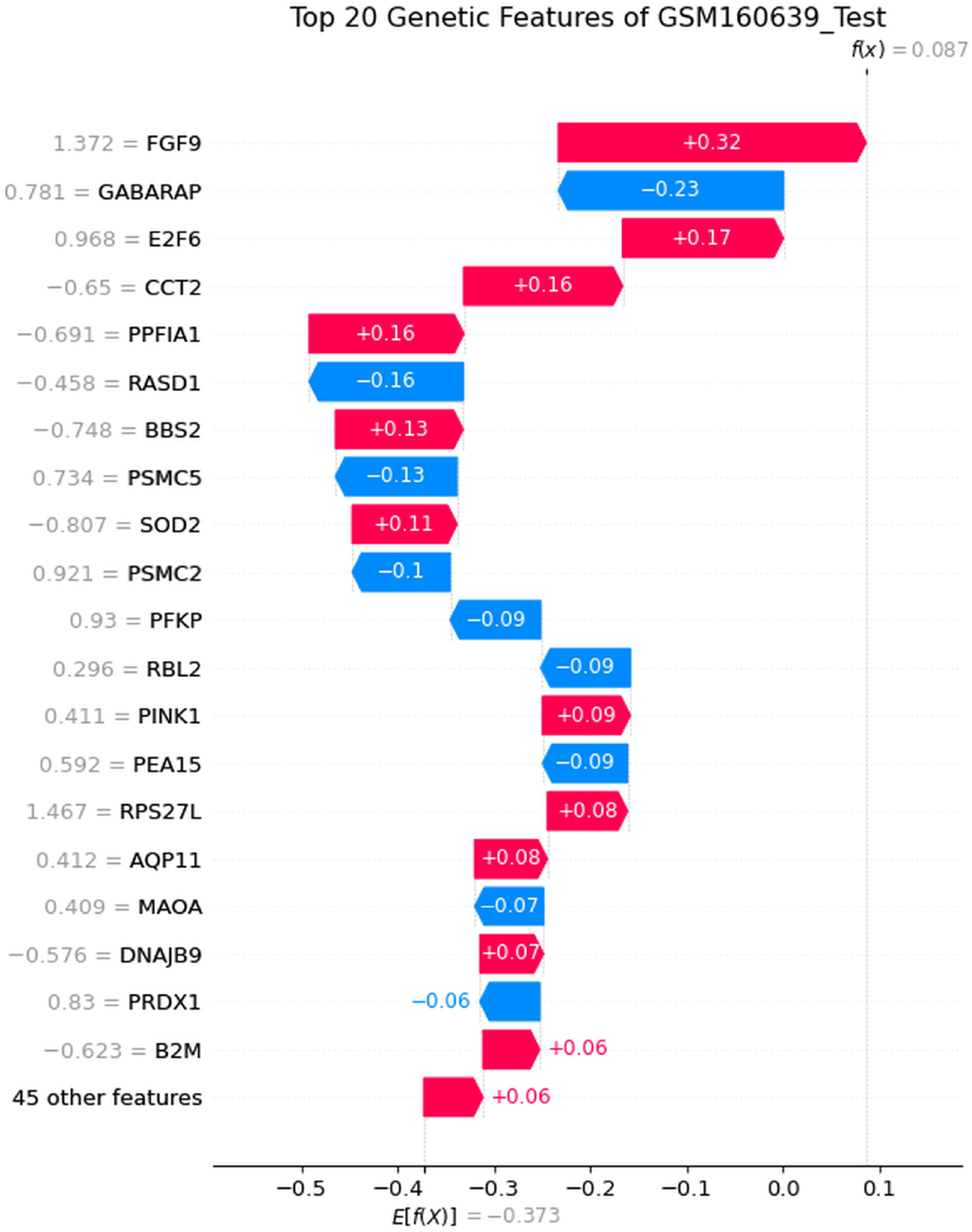
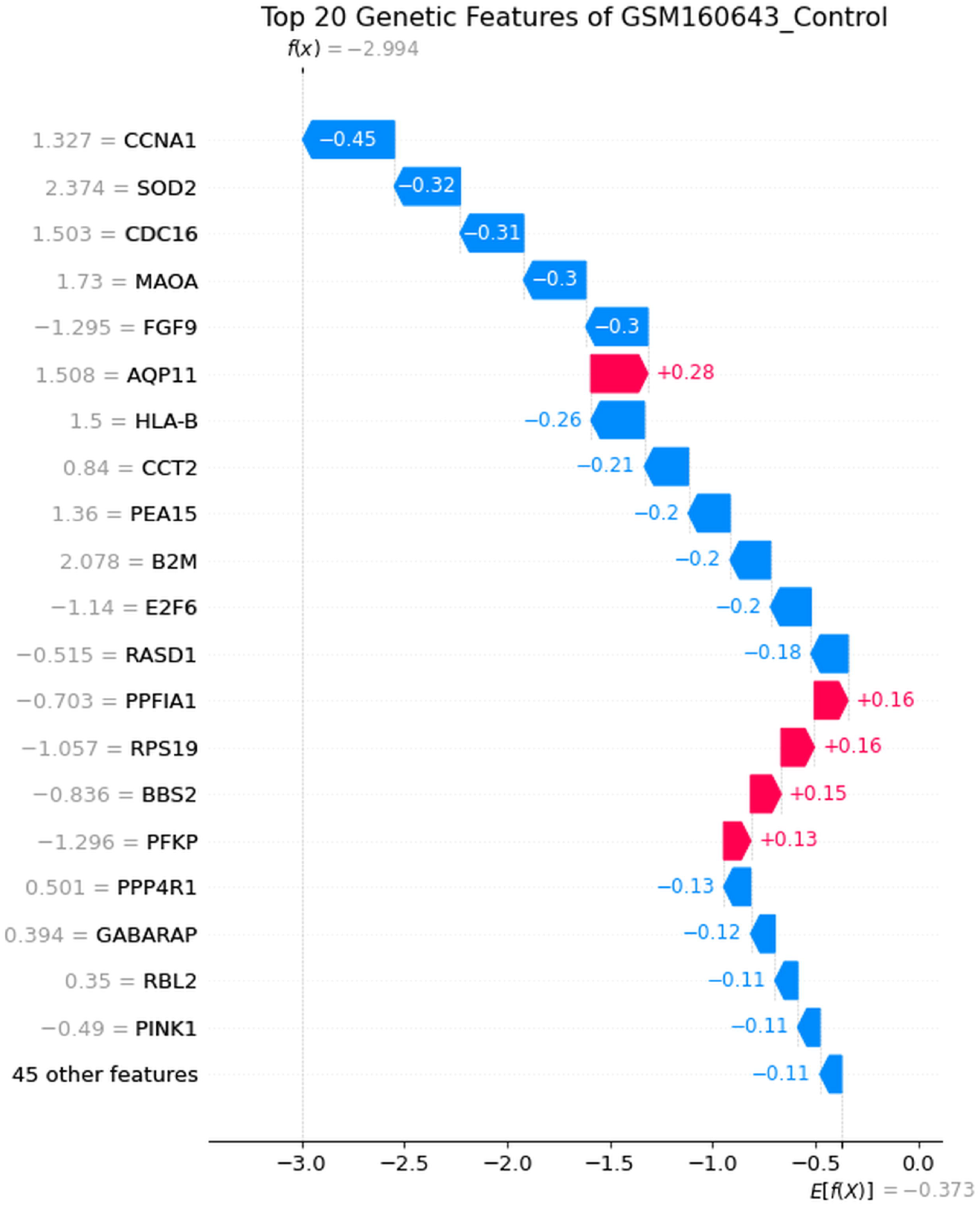
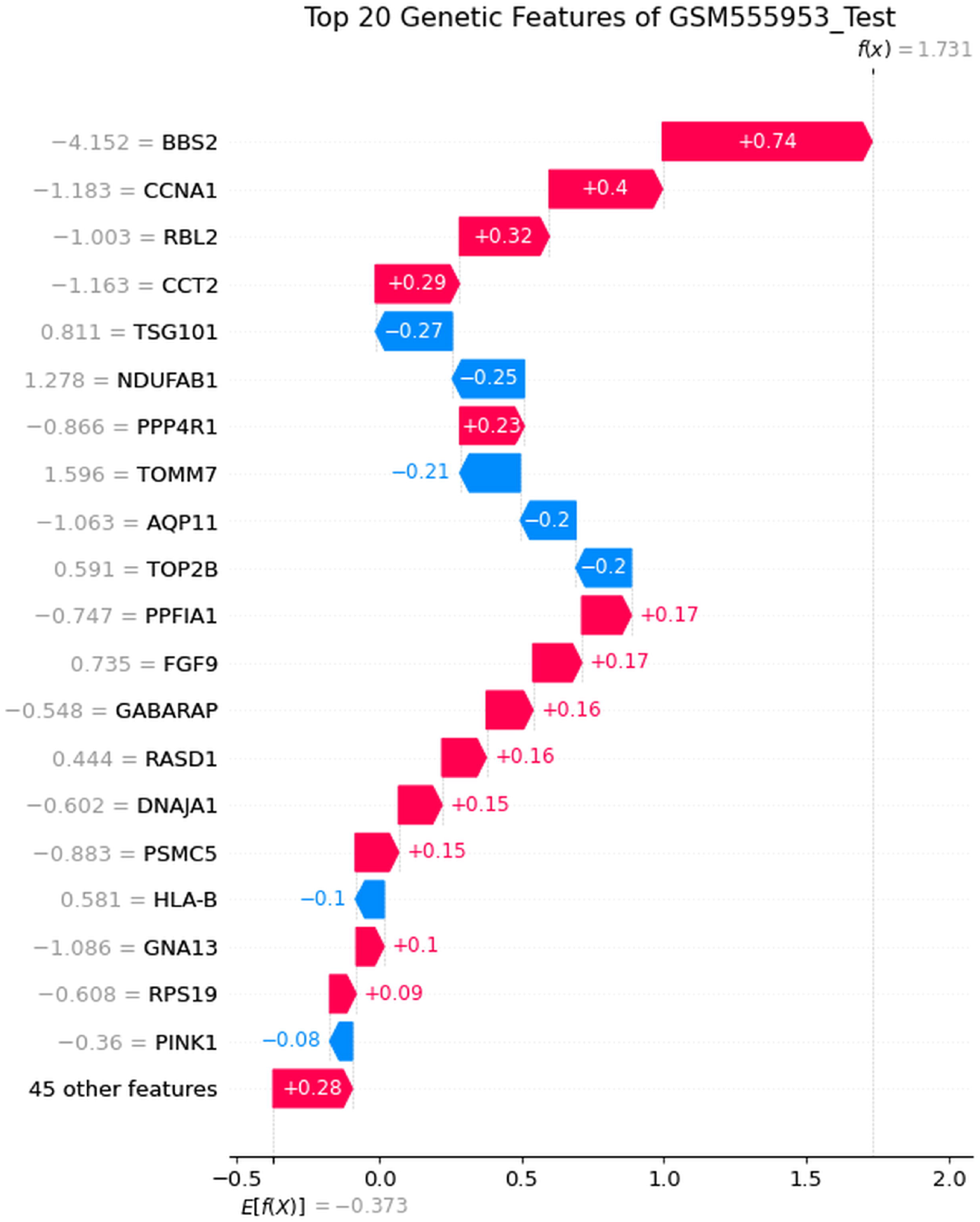
Waterfall plot of representing the contribution of top 20 genes. Waterfall plot depicts the contribution of top 20 genes sorted in the order of their importance based on SHAP value corresponding to (a) GSM160639_Test, (b) GSM160643_Control and (c) GSM555953_Test samples Here x-axis represents the SHAP value with E[f(x)]=-0.373 as the base value or expected value of SHAP explainer and y-axis represents the gene names

The waterfall diagram representing the expected value of the SHAP explainer calculated over the training dataset as -0.373, denoted by φ_0_ in ***equation 1***. The SHAP value corresponding to the GSM160639_Test, GSM160643_Control and GSM555953_Test are 0.087, -2.994 and 1.731 respectively. Furthermore, if we put this shap values in ***equation 2***, we can obtain the prediction probabilities as shown in ***Table 3***. Thus, we can conclude that our model can provide a prediction probability regarding the fertility status of a sperm sample with specific contribution of the feature gene set in support of such prediction.

In order to test the robustness of our model, we used asthenoteratozoospermia (a condition where there is defect in sperm motility as well as morphology) samples viz. GSM4878477 and GSM4878478 as unknown test samples to predict the outcome. Our model correctly classified these samples to be infertile with a probability of 0.99 in both the cases.

## DISCUSSION AND CONCLUSION

The two major causes of male fertility defects pertain to asthenozoospermia and teratozoospermia where there is reduced sperm motility of < 40% or progressive motility with <32% [29] and spermatozoa with abnormal morphology over 85% [30]. In this work we have tried to unveil the genetic reasons underlying the aberrant sperm population. This has been further used to develop a prediction model named **E-InfertilityTest** to test and classify an unknown sperm sample as fertile or infertile. However, the current computational framework does not support classification into infertility subtypes, as it would compromise generalizability of the model with limited data.

Apart from classification of a sperm sample as fertile or infertile, **E-InfertilityTest** will also render the genetic basis behind such classification. Thus it will provide a molecular insight into the causality behind such infertility in a case-to-case basis. However, due to sparsity in the human asthenozoospermia (immotile sperm) and teratozoospermia (aberration in sperm morphology) transcriptome dataset in public domain, the Training Dataset is relatively small. With the availability of more data in future, an increase in the size of the training dataset will yield better result.

It is important to mention that the mature sperm transcriptome bear the ‘transcriptional legacy’ or the molecular archive of the gene expression events that occurred during spermatogenesis [31] on one hand. On the other hand, there is presence of diverse RNA populations corresponding to functionally relevant molecules in mature sperm, other than mere residual remnants from spermatogenesis [32] that may influence sperm fertility profile. The detailed mode of regulation of a significant set of genes (comprising the ML feature set) influence various events which effectively regulates sperm motility and morphology (as depicted in **Figure 7**).

**Figure 7:**
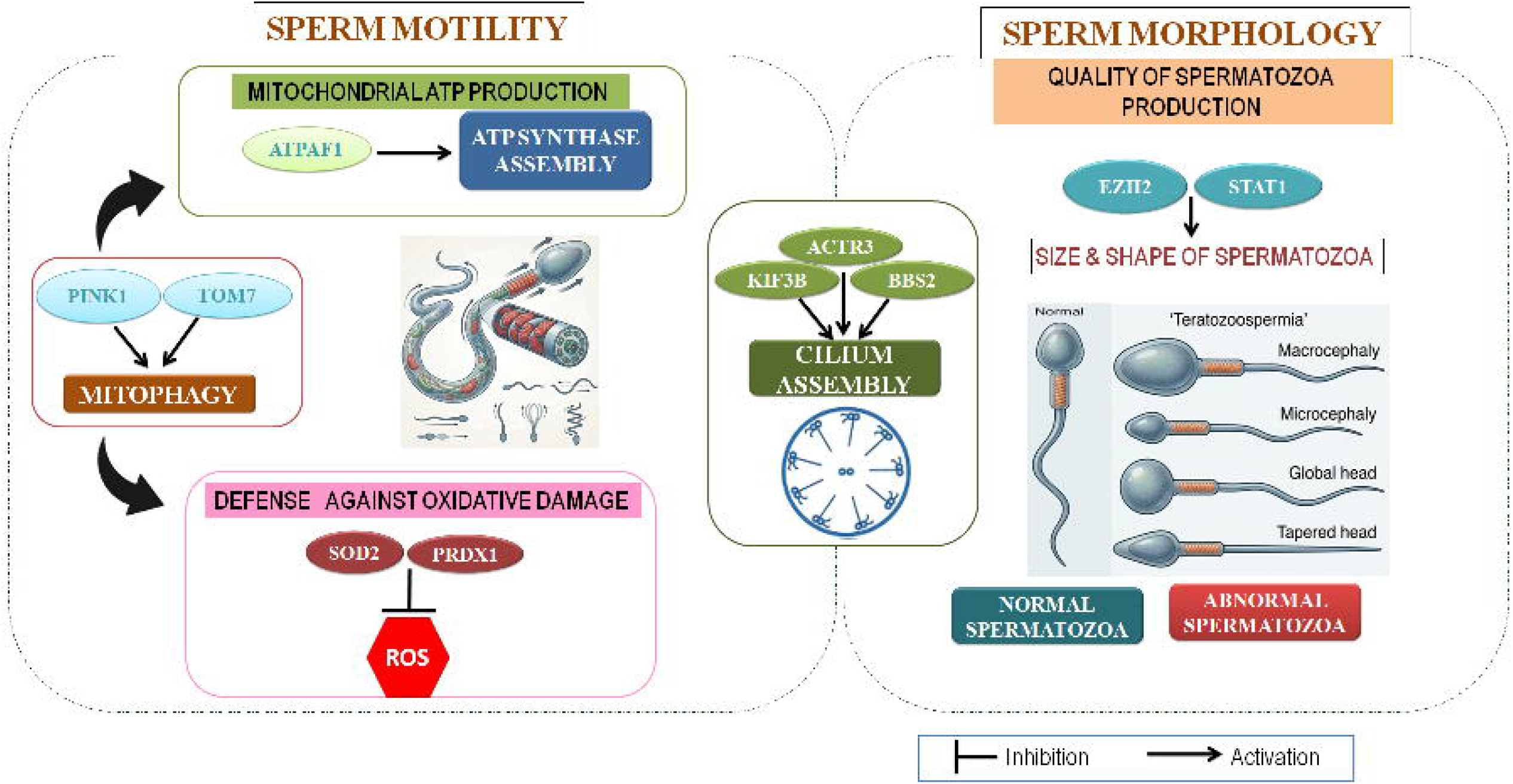
Schematic diagram depicting the regulatory pathways and the associated genes modulating male infertility.

### Sperm Motility

Mitochondrion ATP is the main energy producer behind the **sperm motility** [33]. Mitochondrial oxidative phosphorylation gets compromised due to disruption of ATP synthase assembly caused due to downregulation of ATPAF1 [34]. This reduces sperm movement by reducing ATP production. PINK1 is a mitophagy related gene, which with the help of TOMM7 anchors within the mitochondria, which enables mitophagy [35] to proceed. Downregulation of both of these prevent mitophagy and causes accumulation of abnormal mitochondria, resulting in oxidative stress, reduced ATP production and ultimately affect sperm motility [36]. Reduced expression of SOD2 also results in enhancement of ROS affecting sperm motility [37]. PRDX1 gene encodes Peroxiredoxin-1 (PRDX1) which also protects the cells from oxidative damage. Downregulation of PRDX1 gene possibly produces lower amount of this antioxidant enzyme which in turn is associated with impaired sperm function as reported by [38, 39]. Low levels of Peroxiredoxins have an implication on asthenozoospermic infertile men [40].

### Sperm Morphology

Interestingly, the transcription factor EZH2 has been reported to be downregulated in teratozoospermia [41]. Its deletion resulted in reduced and abnormal spermatozoa production [42]. Further, downregulated STAT1 plays a significant role towards inducing teratozoospermia [43].

Another important event that effects both sperm motility and morphology is cilium assembly (ciliogenesis) where axoneme plays a crucial role. KIF3B regulates intraflagellar transport that delivers axonemal components during ciliogenesis [44]. Its gene variant leads to sperm morphology and motility defects [45]. Another gene BBS2 is important for ciliary membrane trafficking to maintain proper functioning of axoneme [46]. It also affects sperm motility and morphology [47]. Downregulation of KIF3B and BBS2 hampers cilium assembly. ACTR3 gene codes for actin associated protein which helps in ciliary organization [48]. Downregulated ACTR3 cannot maintain proper junction architecture and actin polymerization [49] thus contributing towards sertoli cell dysfunction lowering sperm motility.

Thus, **E-InfertilityTest** is capable of classifying between fertile versus infertile sperm sample especially in “borderline” or idiopathic cases where phenotype alone does not offer clear explanation. Further, it will provide patient specific dominant gene expression profile responsible for the aberration. Future validation experiments incorporating patient samples will substantiate the model performance in a clinical setting. Moreover, this explainable prediction model, once refined with larger dataset, will increase the model’s robustness and may lead towards a more comprehensive understanding of the genetic factors influencing male infertility.

## Supporting information

https://bicresources.jcbose.ac.in/zhumur/download_supple/

## DATA AVAILABILITY

The detail information about the analyzed dataset can be extracted from **Table 1 & Table 2** and the associated dataset can be downloaded from NCBI GEO website (https://www.ncbi.nlm.nih.gov/gds).

## ACKNOWLEDGEMENTS

This work is funded by **National Network project of Bose Institute with Indian Statistical Institute and Vidyasagar University**, sanctioned by the Department of Biotechnology, Government of India, vide the sanction no. BT/PR40176/BTIS/137/84/2023 as well as Indian Council of Medical Research (ICMR) (sanction no. RBMH/FW/2020/10), Government of India.

## AUTHOR CONTRIBUTION

ZG hypothesized, conceptualized and provided overall supervision of the study. ZG and GD initially designed the experiments. Data curation and analysis was done by GD and BG. GD, BG and ZG drafted the manuscript.

---

**Supplementary File S1**: Pathway analysis corresponding to the 65 genes constituting the ML feature set

